# Epigenome-wide Analysis Identifies Genes and Pathways Linked to Neurobehavioral Variation in Preterm Infants

**DOI:** 10.1101/445130

**Authors:** Todd M. Everson, Carmen J. Marsit, T. Michael O’Shea, Amber Burt, Karen Hermetz, Steven L Pastyrnak, Charles R Neal, Brian S Carter, Jennifer Helderman, Elisabeth C. McGowan, Lynne M Smith, Antoine Soliman, Julie A Hofheimer, Sheri A DellaGrotta, Lynne M Dansereau, James F Padbury, Barry M Lester

## Abstract

**Background & Objectives:** Neonatal neurobehavioral performance measures, such as the NICU Network Neurobehavioral Scale (NNNS), have been developed to assess the neurobehavioral characteristics of infants and provide insights into future developmental trajectories. The identification of molecular biomarkers of very early life neurobehavioral experiences could lead to better predictions of the long-term developmental outcomes of high-risk infants including preterm infants. To this end, we aimed to examine whether variability in DNA methylation (DNAm) or epigenetic age from surrogate tissues are associated with NNNS profiles in a cohort of infants born less than 30 weeks postmenstrual age (PMA).

**Methods:** This study was performed within the Neonatal Neurobehavior and Outcomes in Very Preterm Infants (NOVI) Study and included those infants with complete NNNS assessment data and DNAm measured from buccal cells, collected at near term-equivalent age using the Illumina EPIC array (N=536). We tested whether epigenetic age and age acceleration differed between infants based on their NNNS profile classifications. Then we performed an epigenome-wide association study, to test whether DNAm at individual epigenetic loci varied between these NNNS profile groupings. Models were adjusted for recruitment site, infant sex, postmenstrual age, and estimated tissue heterogeneity.

**Results:** We found that infants with an optimal NNNS profile had slightly older epigenetic age than other NOVI infants (β_1_ = 0.201, p-value = 0.026), and that infants with an atypical NNNS profile had differential methylation at 29 CpG sites (FDR < 10%). The genes annotated to these differentially methylated CpGs included *PLA2G4E*, *TRIM9*, *GRIK3*, and *MACROD2*, which have previously been associated with neurological structure and function, or with neurobehavioral disorders.

**Conclusions:** Greater epigenetic age is associated with optimal NNNS responses while altered DNAm of multiple genes are associated with an atypical neurobehavioral profile at near-term equivalent age. These findings build upon the existing evidence that epigenetic variations in buccal cells may serve as markers of neonatal neurobehavior and might facilitate early identification of children at risk for abnormal developmental outcome.

## INTRODUCTION

Preterm birth is a significant global public health problem. In the United States one in eight children are born less than 37 weeks of gestation (1). Survival of infants born extremely preterm, less than 30 week postmenstrual age (PMA), has improved due to technological and medical advancements (1-3). Nonetheless, these infants have increased risk for morbidity and mortality. Children born extremely preterm term are more likely to suffer from chronic illnesses, potentially devastating brain injuries, and adverse neuromotor, cognitive, and behavioral outcomes that persist through adulthood (4-15). These serious health consequences require extensive healthcare, educational and psychosocial community resources, in addition to increased burden on the families and caregivers of these children, emotionally and financially.

It is important to note that not all preterm infants will experience the same health consequences. In the neonatal intensive care unit (NICU), there are limited assessments capable of predicting which infants will experience adverse neurodevelopmental or health outcomes. Identifying those at the greatest risk at the earliest postnatal age maximizes the benefits of interventions aimed at ameliorating long term functional deficits. There is growing evidence that neonatal neurobehavior as measured by the NICU Network Neurobehavioral Scale (NNNS) (16) predicts developmental deficits in infants born preterm and others at risk, beyond what can be predicted based on the assessment of medical risk factors throughout the newborn’s hospital stay (17-20). Poorer performance on the NNNS has also been shown to be predictive of non-optimal developmental outcomes through early childhood (21). Beyond the neurobehavioral and medical assessments, molecular biomarkers may provide insights into how the environment and experiences of the preterm newborn are internalized and may hold additional value as predictive tools useful in risk stratification.

Epigenetics refers to mitotically and meiotically heritable changes in gene expression potential that are not explained by changes in DNA sequence. The most thoroughly studied epigenetic mechanism is DNA methylation (DNAm), particularly in the context of cytosine-phosphate-guanine (CpG) motifs and islands. These methylation marks can be inherited, established in-utero and/or affected by the environment throughout life, thus representing a truly integrated measure of exposure and disease susceptibility. In preterm infants, variability in (DNAm) of candidate genes have been related to medical complications such as sepsis (22), pain related stress (23, 24), medical and neurobehavioral risk (25, 26), and as a potential moderator of NICU environment stress on serotonergic tone and temperament (27). We have also used an epigenome-wide scan of (DNAm) in the placenta to demonstrate relationships between methylation of the *FHIT* and *ANDKR11* genes, which had been previously linked to neurodevelopmental and behavioral outcomes, and performance on the NNNS attention scale in a cohort of term newborns (28).

In order to develop models incorporating molecular biomarkers with performance measures to predict long-term health in preterm infants, it is important to demonstrate whether variability in (DNAm) in accessible tissues is associated with behavioral measures and is a potentially useful indicator of newborn experience. In this study, we hypothesized that infants with either atypical or optimal neurobehavioral profile on the NNNS will exhibit unique patterns of (DNAm). In a U.S. multisite cohort of infants born less than 30 weeks PMA, we profiled genome-wide (DNAm) from buccal swab samples using the Illumina MethylationEPIC array platform. We then tested for differences in epigenetic age and in (DNAm) among infants with optimal or atypical neurobehavioral profiles, which were determined via NNNS latent profile classification.

## METHODS

### Study Population

The Neonatal Neurobehavior and Outcomes in Very Preterm Infants (NOVI) Study was conducted at 9 university-affiliated NICUs in Providence, RI, Grand Rapids, MI, Kansas City, MO, Honolulu, HI, Winston-Salem, NC, and Torrance and Long Beach CA from April 2014 through May 2016. Enrollment and consent procedures for this study were approved by the institutional review boards of all participating institutions. Eligibility was determined based on the following inclusion criteria: 1) birth at <30 weeks’ gestation; 2) parental ability to read and speak English, Spanish, Japanese, or Chinese; 3) residence within 3 hours of the NICU and follow-up clinic. Infants were excluded if their medical record indicated presence of a major congenital anomaly, defined as being in a list of anomalies used in studies from the Eunice Kennedy Shriver National Institute of Child Health and Human Development Neonatal Research Network. These included central nervous system, cardiovascular, gastrointestinal, genitourinary, chromosomal, and nonspecific anomalies. (29, 30) Parents of eligible infants were invited to participate in the study at 31-32 weeks PMA, or when survival to discharge was determined to be likely by the attending neonatologist. Demographic variables, including infant gender, race, ethnicity, maternal education and partner status were collected from the maternal interview. Medical variables, including birthweight, gestational age, length of stay, weight at discharge, and gestational age at discharge were abstracted from the medical records. Overall, 665 infants were enrolled, from whom complete neurobehavioral and medical data were obtained on 647, and buccal cells were collected on 624 of these for infants for epigenetic screening.

### NICU Network Neurobehavioral Scale (NNNS)

Neonatal neurobehavior was assessed using the NNNS. The NNNS is a 20-30 minute standardized procedure that includes measures of active and passive tone, primitive reflexes, items that reflect physical maturity, social and behavioral functioning including visual and auditory tracking, cuddling and soothability, and a checklist of stress signs organized by organ system (31). The NNNS was administered at 36 +/- 2 weeks PMA or during the week of NICU discharge by site examiners who were trained and certified by central NOVI NNNS trainers. The exam was conducted 45 minutes prior to a scheduled feeding or routine care in order to maximize alertness and avoid disrupting NICU routines that facilitate sleep patterns. Individual items were converted to 12 summary scores: attention, handling, self-regulation, arousal, excitability, lethargy, hypertoniicity, hypotonicity, non-optimal reflexes, asymmetric reflexes, quality of movement and stress abstinence. Summary scores were converted to NNNS profiles, which are mutually exclusive, discrete categories representing the infant’s pattern of performance across the summary scores (21).

### DNA Methylation (DNAm) Analysis

Genomic DNA was extracted from buccal swab samples, collected near term-equivalent age, using the Isohelix Buccal Swab system (Boca Scientific), quantified using the Quibit Fluorometer (Thermo Fisher, Waltham, MA, USA) and aliquoted into a standardized concentration for subsequent analyses. DNA samples were plated randomly across 96-well plates and provided to the Emory University Integrated Genomics Core for bisulfite modification using the EZ DNA Methylation Kit (Zymo Research, Irvine, CA), and subsequent assessment of genome-wide (DNAm) using the Illumina MethylationEPIC Beadarray (Illumina, San Diego, CA) following standardized methods based on the manufacturer’s protocol. Samples with more than 5% of probes yielding detection p-values > 1.0E-5 (74 samples), with mismatch between reported and predicted sex (7 samples), or incomplete covariate data (7 samples) were excluded. Additionally, probes with median detection p-values < 0.05 were excluded. Array data were normalized via functional normalization, then standardized across Type-I and Type-II probe designs with beta-mixture quantile normalization (32). Probes that measured methylation on the X and Y chromosomes, probes that had single nucleotide polymorphisms (SNP) within the binding region, that could cross-hybridize to other regions of the genome (33), or probes that had low variability (range of beta-values < 0.05) (34) were excluded. After exclusions, 690,781 probes were available from 536 samples for this study. We used gaphunter to flag probes that had outliers or distributional issues that may be related to genetic effects on (DNAm) measurement (35).

### Estimate of Epigenetic Age

We estimated epigenetic age using the online (https://labs.genetics.ucla.edu/horvath/dnamage/) epigenetic clock calculator (36). This method utilizes (DNAm) levels at previously identified CpGs that are predictive of chronological age and has been shown to be highly accurate across a wide range of different cell and tissue types (37). This clock also calculates two measures of age acceleration: the difference between epigenetic and chronological age, and the residuals when epigenetic age is regressed on chronological age in a linear model. We investigated the age acceleration residuals and epigenetic age in this study.

### Estimates of Tissue Heterogeneity

DNAm differs between cell-types, and cellular heterogeneity presents a likely source of confounding in epigenome-wide association studies of mixed cell samples (38). Thus, we estimated the proportions of epithelial, fibroblast, and immune cells in our cheek swab samples using reference methylomes (39) and adjusted for cellular heterogeneity by including the proportions of cells types as covariates in the statistical models.

### Statistical Analyses

Latent profile analysis (LPA, Mplus version 8.1) was used to group infants into mutually exclusive categories using 12 NNNS summary scores based on previous work (21). Membership in categorical latent profiles that represent heterogeneous subgroups was inferred from the 12 NNNS variables. LPA models with different numbers of profiles were fitted. Determination of the best model fit was assessed via Bayesian information criteria (BIC) adjusted for sample size, whereby the smallest BIC value indicates the best fit as well as minimization of cross classification probabilities.

Statistical analyses of epigenomic data were performed in R version 3.5. We tested for differences in epigenetic age and age acceleration between the atypical NNNS profile and optimal NNNS profile versus those in the other NNNS profiles using robust linear models via the MASS package. Standard errors and p-values for robust regressions were estimated using White’s sandwich estimator to protect against potential heteroscedasticity. Epigenetic age and age acceleration were included as continuous dependent variables, the NNNS profiles were included as a three-level factor for the independent variable, while adjusting for sex, recruitment site, and cellular heterogeneity. The epigenome-wide association study was performed with robust linear regressions for each CpG site that passed QC, regressing methylation beta-values on the NNNS profiles, while adjusting for sex, PMA, and proportions of epithelial cells and fibroblasts. QQ-plots and Manhattan plots were produced using the qqman package. To account for multiple testing, we implemented a false discovery rate (FDR) and considered those associations that were within a 10% FDR to be statistically significant. We report all results from models that yielded suggestive associations (p-value < 0.0001) in the supplemental materials.

To gain insights into the biological functions of the NNNS-associated CpG sites, we performed over-representation analyses with ConsensuPathDB (CPDB) (40, 41). We utilized CPDB to examine our gene lists for enrichment with neighborhood-based entity sets (NESTs) with a radius of one, pathway-based gene sets from KEGG, Biocarta, and Reactome with minimum overlap with our gene-set of 2, and gene-ontology (G0) terms. Over-representation results within a 10% FDR were determined to be statistically significant. For over-representation analyses, we utilized a gene-list containing the genes annotated to the top 250 CpG sites from the EWAS that were associated with the atypical NNNS profile. We also aimed to examine whether our NNNS-associated CpGs were within genes that have been linked with phenotypes related to neurodevelopment or neurodegeneration. Thus, we annotated the top 250 CpGs with traits that have been linked to genes via the genome-wide association study database (GWASdb) (42).

## RESULTS

### Study Population and NNNS Profile Results

We identified six distinct NNNS profiles representing groups of infants with similar neurobehavioral responses (Table 1). Two of these profiles stood out as particularly distinctive (Figure 1). Infants in Profile 1 had the most optimal performance with the best attention and regulations scores, an average requirement for handling, typical motor movement and few signs of stress. Infants in Profile 6 showed atypical performance with poor attention, a substantial requirement for handling, poor regulation, exceptionally high arousal and excitability, hypertonia, poor quality of movement and substantial signs of stress. Thus, Profile 1 represents an optimal profile characterized by generally positive neurobehavioral responses, while Profile 6 represents a infants with atypical neurobehavioral responses. These findings are similar to profiles observed previously by others (21) and in our own research (26). To limit the number of tests being performed, the current study focused on the optimal (Profile 1) and atypical (Profile 6) profiles, while using the combination of Profiles 2-5 as the referent category in downstream analyses. Average PMA at birth, PMA at buccal cell collection, and maternal age did not substantially differ between the different NNNS profile groupings (Table 2). On the other hand, we did find that a larger proportion of infants with atypical profiles had caregivers with lower socioeconomic status (SES) (16.7%) and lower educational attainment (22.2%), compared to those in the optimal group (4.8% and 6.5% respectively). We also observed substantial differences in NNNS profile assignment by recruitment site, and thus recruitment site was controlled for in all downstream analyses.

**Figure 1:**
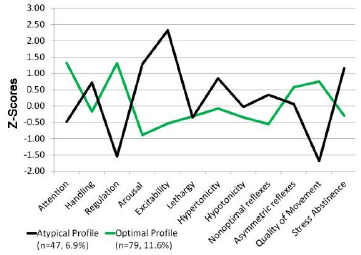
NNNS z-scores across individual assessments for the atypical profile (black) and the optimal profile (green) among all NOVI infants assessed for the NNNS ().

**Table 1:** Means and standard errors of individual NNNS assessment scores across the NNNS profile groupings identified by LPA (N=536).

| Assessment | Profiles |  |  |  |  |  | ANOVA<br>p-val. |
| --- | --- | --- | --- | --- | --- | --- | --- |
|  | 1<br>(n=62) | 2<br>(n=176) | 3<br>(n=61) | 4<br>(n=83) | 5<br>(n=118) | 6<br>(n=36) |  |
| Attention | 7.27<br>(1.14) | 5.04<br>(1.22) | 4.89<br>(0.90) | 6.04<br>(1.18) | 4.43<br>(1.36) | 4.36<br>(1.25) | < 0.001 |
| Handling | 0.37<br>(0.25) | 0.29<br>(0.22) | 0.61<br>(0.25) | 0.55<br>(0.22) | 0.37<br>(0.26) | 0.62<br>(0.27) | < 0.001 |
| Quality of Movement | 5.06<br>(0.55) | 4.89<br>(0.45) | 4.9<br>(0.59) | 4.58<br>(0.58) | 4.23<br>(0.49) | 3.4<br>(0.64) | < 0.001 |
| Self Regulation | 6.69<br>(0.60) | 5.82<br>(0.45) | 5.04<br>(0.53) | 6.09<br>(0.58) | 5.21<br>(0.51) | 4.44<br>(0.72) | < 0.001 |
| Non-Optimal Reflexes | 4.18<br>(1.61) | 4.83<br>(1.52) | 4.84<br>(1.67) | 4.1<br>(1.50) | 7.47<br>(1.70) | 6.22<br>(1.93) | < 0.001 |
| Stress Abstinence | 0.11<br>(0.06) | 0.09<br>(0.06) | 0.14<br>(0.07) | 0.17<br>(0.06) | 0.16<br>(0.06) | 0.22<br>(0.08) | < 0.001 |
| Arousal | 3.18<br>(0.48) | 3.55<br>(0.37) | 4.59<br>(0.51) | 3.91<br>(0.50) | 3.47<br>(0.50) | 4.61<br>(0.51) | < 0.001 |
| Hypertonicity | 0.34<br>(0.75) | 0.27<br>(0.55) | 0.8<br>(1.15) | 0.25<br>(0.49) | 0.23<br>(0.53) | 0.97<br>(1.18) | < 0.001 |
| Hypotonicity | 0.05<br>(0.22) | 0.15<br>(0.36) | 0.13<br>(0.39) | 0.06<br>(0.29) | 0.52<br>(0.72) | 0.25<br>(0.44) | < 0.001 |
| Asymmetrical Reflexes | 1.69<br>(1.52) | 0.43<br>(0.73) | 0.25<br>(0.54) | 2.05<br>(1.51) | 0.94<br>(1.15) | 0.97<br>(1.44) | < 0.001 |
| Excitability | 1.42<br>(1.15) | 1.16<br>(0.93) | 4.77<br>(1.27) | 2.52<br>(1.35) | 2.18<br>(1.13) | 7.22<br>(1.61) | < 0.001 |
| Lethargy | 3.77<br>(1.58) | 4.8<br>(1.87) | 3.64<br>(1.53) | 3.17<br>(1.58) | 6.17<br>(2.15) | 3.97<br>(1.92) | < 0.001 |

ANOVA: analysis of variance, LPA: latent profile analysis, NNNS: NICU network neurobehavioral scale.

**Table 2:** Distributions of covariates by NNNS profile groupings.

| Sample Characteristics | Profiles 2,3,4,5<br>(N=438) | Optimal<br>(N=62) | Atypical<br>(N=36) |
| --- | --- | --- | --- |
| Infant Gender: Male | 55.3% (242/438) | 56.5% (35/62) | 61.1% (22/36) |
| Recruitment Site: |  |  |  |
| WIH | 19.4% (85/438) | 3.2% (2/62) | 19.4% (7/36) |
| SHD | 22.6% (99/438) | 1.6% (1/62) | 0.0% (0/36) |
| KMC | 16.0% (70/438) | 17.7% (11/62) | 19.4% (7/36) |
| CMH | 14.8% (65/438) | 8.1% (5/62) | 0.0% (0/36) |
| WFU | 14.6% (64/438) | 67.7% (42/62) | 22.2% (8/36) |
| LAB | 12.6% (55/438) | 1.6% (1/62) | 38.9% (14/36) |
| PMA at Buccal Collection (weeks) | 38.930 ± 3.150 | 39.415 ± 4.012 | 40.210 ± 3.462 |
| PMA at Birth (weeks) | 27.091 ± 1.881 | 26.756 ± 1.954 | 26.452 ± 2.154 |
| Birth Weight (grams) | 969.0 ± 281.4 | 903.8 ± 265.0 | 861.9 ± 291.4 |
| Maternal Age at childbirth (years) | 29.171 ± 6.297 | 29.095 ± 6.972 | 27.972 ± 6.396 |
| Education: less than High School/GED | 14.6% (62/426) | 6.5% (4/62) | 22.2% (8/36) |
| Low SES | 8.2% (35/427) | 4.8% (3/62) | 16.7% (6/36) |

PMA: Postmenstrual Age, SES: socioeconomic status, GED: General Equivalency Diploma, NNNs: NICU network neurobehavioral scale,

### Epigenetic Age and Age Acceleration Associations with NNNS Profiles

Epigenetic age negatively correlated with gestational age at birth (R^2^ = 0.06, p-value < 0.0001), but positively correlated with age since birth (R^2^ = 0.15, p-value < 0.0001) and PMA (R^2^ = 0.14, p-value < 0.0001) (Supplemental Figure 1). We examined differences in epigenetic age and age acceleration that associated with the optimal (n=62) and atypical (n=36) NNNS profiles, by comparing them to the rest of the NOVI infants (n=438). We found that the infants in the optimal profile tended to have significantly older epigenetic age (β_1_ = 0.201, p-value = 0.026) whereas that atypical profile exhibited no difference in epigenetic age (β_1_ = −0.022, p-value = 0.84) when compared to the rest of the NOVI infants (Figure 2A) age acceleration did not significantly differ when comparing the optimal or atypical profiles to the rest of the NOVI infants, though we did observe a step-wise pattern in which the atypical profile had the lowest estimated age acceleration (β_1_ = −0.121, p-value = 0.22) and the optimal profile had the greatest estimated age acceleration (β_1_ = 0.082, p-value = 0.36) (Figure 2B). These models were adjusted for sex, recruitment site, postmenstrual age, and estimated proportions of epithelial cells and fibroblast cells.

**Figure 2:**
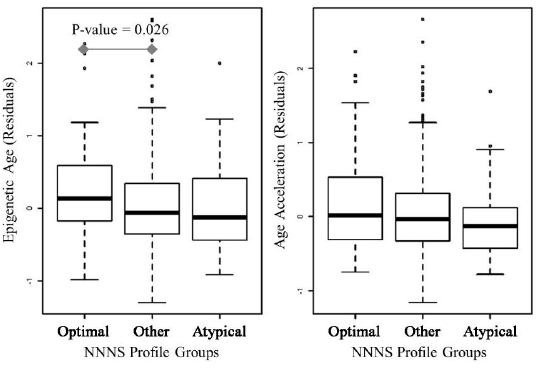
Relationships between epigenetic age (A) and age acceleration (B) with NNNS profile groupings; the x-axis includes the optimal profile (Profile 6), the atypical profile (Profile 1), as well as other NOVI infants (Profiles 2-5), while the y-axis represents the residuals of epigenetic age and age acceleration after adjusting for sex, recruitment site, and cell-mixture.

**Figure 3:**
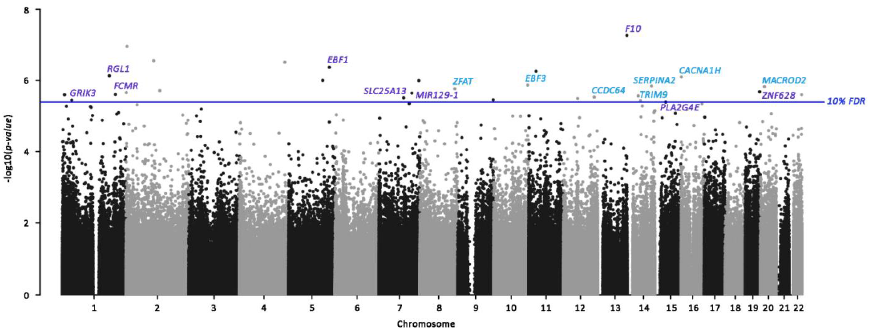
Manhattan plot of epigenetic loci associated with the atypical NNNS profile; the x-axis represents the genomic location of the individual probes and the y-axis represents the ‐log_10_(p-values) from related to the atypical NNNS profile, adjusted for sex, recruitment site, postmenstrual age, and cell-mixture; gene annotations for the CpGs yielding associations within the 10% FDR threshold have been added to the plot.

### Epigenome-Wide Association Study of NNNS Profiles

We then performed an EWAS of these two NNNS profiles to examine whether underlying patterns of epigenetic regulation measured in buccal cells differed among infants within the NNNS profiles. We report all results for the EWAS of the optimal and atypical NNNS profiles that yielded associations with p-values < 0.0001 in the Supplemental Materials (Supplemental Table 1 & Supplemental Table 2). We identified 30 CpGs that were differentially methylated (at a 10% FDR) with either of the NNNS profile groupings (Table 3). However, only one of these CpGs associated with the optimal NNNS profile, cg03046148, at which optimal NNNS infants tended to have higher DNAm levels (β_1_ = 0.0145, p-value = 1.43E-07). On the other hand, we identified 29 epigenetic loci that associated with the atypical NNNS profile after FDR-adjustment, which were located throughout the genome (Figure 2). The most statistically significant relationship was observed at cg23172057 (β_1_ = 0.0299, p-value = 5.43E-08) which is within the body of the coagulation factor X (*F10*) gene. The magnitudes of effect among the FDR-significant hits tended to be small, with differential methylation ranging between 0.42% to 6.53% lower and between 0.31% to 10.61% higher methylation among the atypical NNNS group. The CpGs with the largest magnitudes of association among the FDR-significant hits were observed at cg14792155 (β_1_ = 0.1061, p-value = 4.04E-06) which is within the body of the phospholipase A2 group IVE (*PLA2G4E*) gene and at cg07850633 (β_1_ = −0.0653, p-value = 1.49E-06) which is within the body of the MACRO domain containing 2 (*MACROD2*) gene. Because we observed differences in SES among those with optimal or atypical NNNS profiles, we performed a secondary EWAS with additional adjustment for SES, which is strongly correlated with maternal education. These additional adjustments for SES did not alter the observed associations between DNAm and atypical NNNS profiles (Supplemental Figure 2).

**Table 3:**
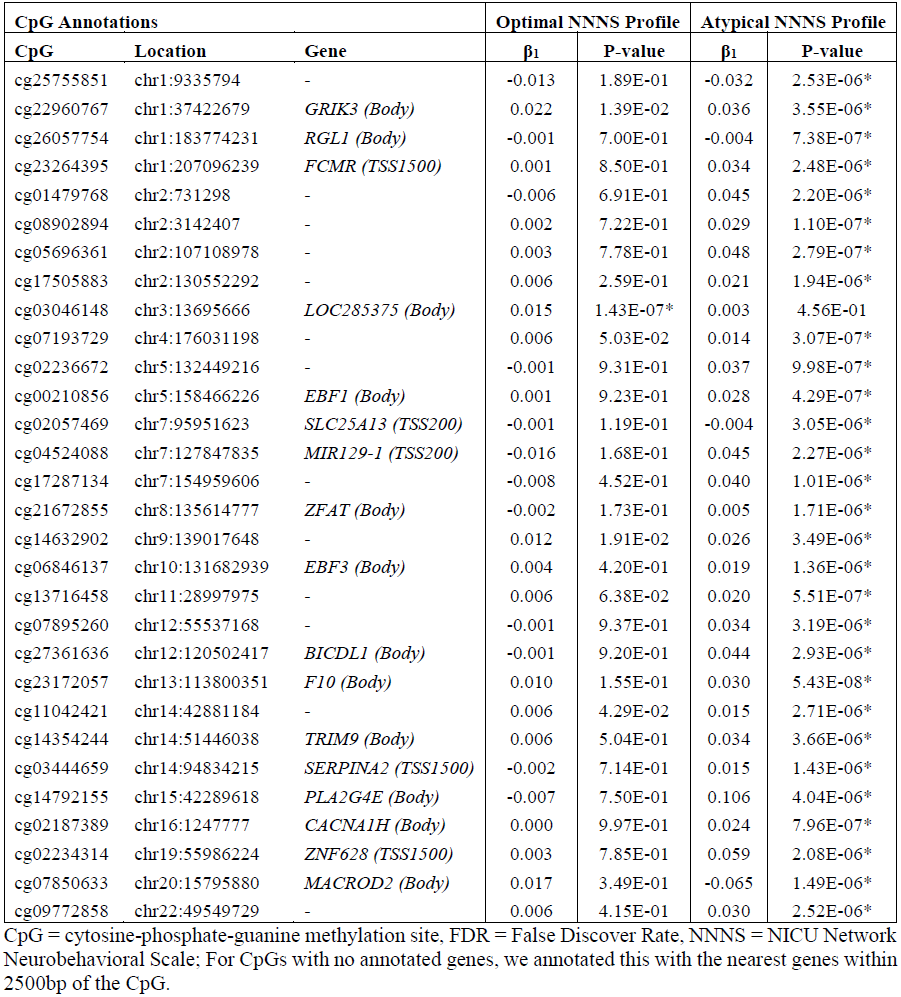
Epigenome-wide association study results for CpG sites that yielded associations within a 10% FDR (*) for either the optimal or atypical NNNS profile groupings; beta coefficients (β_1_) represent the mean difference in methylation proportion when comparing the optimal or Atypical NNNS profiles to the rest of the NOVI sample.

### Functional and Phenotype Enrichment

We then used enrichment analyses to examine whether the genes annotated to NNNS-associated CpGs have a higher than expected proportion of genes that interact with each other, are involved in known biological pathways, or are linked to specific gene ontology terms. For this analysis, we utilized the top 250 CpGs that associated with the atypical NNNS profiles. We found that this gene-set was enriched for one neighborhood-based entity sets (NESTs) (FDR q-value = 0.031), centered on the *CRIM1* gene which has physical interactions with four genes, two of which were also in our gene-set: *ATXN7* and *MEGF6*. We also identified 54 pathway-based gene-sets (Supplemental Table 3) many of which may be relevant for neurodevelopment, including synaptic activity, neurotransmitters, and nerve growth factors (Table 4). Additionally, our gene-set was enriched for nine gene-ontology (G0) terms (Supplemental Table 4), including neuron projection (FDR q-value = 0.0371) and neuron part (FDR q-value = 0.0505) (Table 5). Multiple pathway and GO-term enrichments included *GRIK3*, *TRIM9*, and *PLA2G4E*, genes that were annotated to CpGs that yielded FDR-significant associations from our EWAS.

**Table 4:** Pathways involved in neurodevelopment and/or neuronal activity that were significantly (FDR < 0.10) over-represented among the genes annotated to the top 250 CpGs that associated with the atypical NNNS profile.

| Pathway ID | Pathway Description | Total Genes | NNNS-Associated Genes | P-val. | FDR Q-val. |
| --- | --- | --- | --- | --- | --- |
| path:hsa04724 | Glutamatergic synapse - Homo sapiens (human) | 114 | *PLCB1*; *GRIK3*; *PRKCB*; *PLA2G4E*; *ADCY3* | 0.0020 | 0.026 |
| R-HSA-112314 | Neurotransmitter receptors and postsynaptic signal transmission | 152 | *PLCB1*; *GRIK3*; *AP2A2*; *PRKCB*; *ADCY3* | 0.0069 | 0.040 |
| R-HSA-416993 | Trafficking of GluR2-containing AMPA receptors | 17 | *PRKCB*; *AP2A2* | 0.0077 | 0.040 |
| R-HSA-112316 | Neuronal System | 367 | *PLCB1*; *PRKCB*; *PTPRS*; *KCNK9*; *ADCY3*; *GRIK3*; *AP2A2*; *KCNA3* | 0.0082 | 0.041 |
| ngfpathway | nerve growth factor pathway (ngf) | 18 | *PLCG1*; *NGFR* | 0.0086 | 0.043 |

**Table 5:** Gene-ontology terms that were significantly (FDR < 0.10) over-represented among the genes annotated to the top 250 CpGs that associated with the atypical NNNS profile.

| GO ID | Term | Total Genes | NNNS-Associated Genes | P-val. | FDR Q-val. |
| --- | --- | --- | --- | --- | --- |
| GO:0043005 | neuron projection | 1226 | *PTPRN2; TRIM9; CDH23; PTPRS; IGSF9; TMC2; GRIK3; SAG; MCRS1; PDE6B; PTK2B; MARK4; ATP1A3; CAPN2; RHO; KLC1; MOB2; CLU; KCNA3; RGS12; NGFR* | 5.62E-04 | 0.0371 |
| GO:0097458 | neuron part | 1535 | *IGSF9; TMC2; MCRS1; PTK2B; NGFR; PITX3; KCNA3; TRIM9; KCNK9; SAG; MARK4; CAPN2; RGS12; PTPRN2; CDH23; MOB2; CLU; PDE10A; PTPRS; GRIK3; PDE6B; ATP1A3; RHO; KLC1* | 8.14E-04 | 0.0505 |

### CpG Annotation

Relatively few genes have been studied for their associations with neurobehavioral characteristics shortly after birth. However, it is plausible that the genes that are linked to cognition, neurobehavior, or neurodegeneration at other life stages may also be important in neurobehavioral function in very early life. Thus, we identified phenotypes or traits that have been associated with the genes annotated to the CpGs from our analysis that were associated with NNNS profiles at a 10% FDR (Table 6). Of particular note, 7 of the 11 genes have been linked to neurobehavioral or neurodegenerative traits including autism, attention deficit hyperactivity disorder, cognitive impairment, depression, and psychosis.

**Table 6:** Genes annotated to NNNS-associated CpGs that have been linked to traits from the GWASdb.

| Gene | N | Trait Types | Specific Cognitive, Neurobehavioral or Neurological Traits |
| --- | --- | --- | --- |
| *F10* | 4 | Hematopoietic | - |
| *EBF1* | 34 | Hematopoietic, Cardiovascular, Immune, Growth & Metabolism, Birth Outcomes, Neurobehavior, Cognition | Psychosocial stress measurement, cognitive impairment, cognitive decline measurement |
| *RGL1* | 7 | Neurobehavior, Cardiovascular, Environmental Exposures, Hematopoietic, Liver | Attention deficit hyperactivity disorder, conduct disorder, schizophrenia, response to antipsychotic drug, response to antidepressant |
| *CACNA1H* | 1 | Hematopoietic | - |
| *EBF3* | 3 | Cancer, Bone, Neurological, Sleep | Peripheral neuropathy |
| *SERPINA2* | 6 | Growth, Respiratory, Cardiovascular, Hematopoietic | - |
| *MACROD2* | 31 | Neurobehavior, Neurological, Liver, Growth & Metabolism, Hematopoietic, Bone, Birth Outcomes, Respiratory, Cardiovascular, Environmental Exposures, Bone, Immune | Autism, eating disorder, brain connectivity measurement, sporadic amyotrophic lateral sclerosis, prion disease, mood disorder |
| *ZFAT* | 12 | Growth & Metabolism, Cardiovascular, Cognitive, Environmental Exposures, Immune, Respiratory, Other | Self-reported educational attainment |
| *GRIK3* | 6 | Neurobehavior, Immune, Respiratory, Environmental Exposures | Unipolar depression, neuroticism measurement, depressive symptom |
| *TRIM9* | 5 | Growth & Metabolism, Neurobehavior, Immune, Cancer | Psychosis |
| *PLA2G4E* | 1 | Environmental Exposures | - |

N = number of traits linked to that gene,

## DISCUSSION

Our study focused on a comparison of neurobehavioral profiles with (DNAm) levels and epigenetic age. We used the NNNS summary scores to identify a group of infants with an optimal profile and a group with an atypical profile, which are similar to what has been observed previously by others and in our own research including preterm infants (21, 26, 43). We found that very premature infants in the NOVI cohort with an optimal neurobehavioral profile had older epigenetic age than other very premature infants. We also found that age acceleration followed a stepwise trend in which infants with the optimal profile had the greatest age acceleration and infants with the atypical profile had the least age acceleration, though this finding was not statistically significant. Epigenetic age is an estimate of the state of underlying physiologic processes, as they relate to biological development and maintenance (37), and has been studied in the context of health conditions that are linked to the aging process, including frailty (44), physical capability (45, 46), cognitive fitness (45), decreased cognitive function and neuropathologies in persons suffering from Alzheimer’s Disease (47), and all-cause mortality (48). In these studies, age acceleration is analogous to biological decline. However, in early life, when children are still undergoing substantial growth and development, it is unclear whether accelerated aging would be expected to be associated with positive or negative developmental characteristics. In fact, if epigenetic aging captures, or is a surrogate for, the activity of developmental processes, epigenetic age acceleration throughout early development may be an indicator through which to track developmental “catch-up”. An epigenetic clock has also been developed to estimate epigenetic gestational age acceleration from cord blood (DNAm) (49). Interestingly, gestational age acceleration has been associated with reduced infant respiratory morbidities (50), which provides some evidence that older epigenetic age in infancy may correlate with positive developmental characteristics. However, it is unknown whether early developmental “catch-up” has long-term benefits. This concept may be similar to catch-up growth, in which rapid weight gain in preterm infants and in term growth restricted infants, though beneficial early on, increases the risk of later cardiometabolic outcomes (51), which is consistent with the developmental origins of health and disease paradigm. Studying indicators of developmental catch-up would be particularly important in longitudinal settings and among children born preterm that are underdeveloped at birth and at high risk for adverse developmental outcomes later in childhood.

A handful of studies have examined the relationships between epigenetic age and development in children. For instance, in childhood or adolescence, epigenetic age has been positively associated with measures of physical development such as fat mass, height, Tanner stages (52), pubertal development, internalization and thought problems (53), and increased cortisol (53, 54). One study observed interrelationships between age acceleration, cortisol production, and hippocampal volume, potentially linking hypothalamic-pituitary-adrenal (HPA) axis activity, neuroanatomy, and epigenetic aging (54). Our study provides some evidence that even shortly after birth, older epigenetic age may be an indicator of better neurobehavioral performance. However, this topic requires additional study and should ideally be investigated in a longitudinal manner, in which both epigenetic age and neurobehavioral assessments are tracked in parallel through early-life development.

Though the infants with the atypical NNNS profile did not have significantly different epigenetic age from the other NOVI infants, we did identify multiple differentially methylated CpGs throughout the genome that were associated with the group with atypical neurobehavioral responses. Our EWAS revealed 29 epigenetic loci that significantly associated with the atypical profile and that seven of the genes annotated to these CpGs have been linked to cognition or educational attainment (*EBF1* & *ZFAT*), and neurobehavioral or neurological disorders in GWAS studies (*EBF1*, *RGL1*, *EBF3*, *MACROD2*, *GRIK3*, and *TRIM9*). We also found that the top 250 CpGs from this analysis were enriched for genes within pathways involving neurotransmitters and synaptic activity, as well as GO-terms related to neuron projection and structure. Of particular note, three genes annotated to the FDR-significant CpGs were within these neuronal-associated GO-terms and pathways: phospholipase A2 group IVE (*PLAG2G4E*), glutamate Ionotropic Receptor Kainate Type Subunit 3 (*GRIK3*), and Tripartite Motif Containing 9 (*TRIM9*). The CpG site that yielded the largest magnitude of association among our statistically significant hits, cg14792155, was within the body of the *PLA2G4E* gene. This gene encodes for a calcium-dependent N-acyltransferase; experimental mouse models have implicated that it likely plays a critical role in endocannabinoid signaling in the nervous system (55) and thus differential regulation of this gene has implications for neurodevelopment and neurodegenerative disorders (56). In human observational studies, placental CpGs within *PLA2G4E* have been observed to be differentially methylated in association with extremely preterm births (57) and genetic variants within the *PLA2G4E* gene have been implicated as a potential risk factor for neurodevelopmental problems such as panic disorder (58). The protein encoded by *GRIK3* is involved in presynaptic neurotransmission. Deletion of *GRIK3* may contribute to developmental delay (59) while several single nucleotide polymorphisms within *GRIK3* have been associated with schizophrenia (60), obsessive-compulsive disorder (61), and depression (62). The protein encoded by *TRIM9* regulates axon guidance and neural outgrowth (63, 64), while deletion of *TRIM9* has been associated with structural and functional abnormalities and impaired learning and memory in mice (65).

Though many of the genes annotated to the FDR-significant CpGs were not included in the neuronal-associated GO-terms or pathways, some of these genes have also been implicated in neurodevelopmental or neurodegenerative disorders. The CpG site that yielded the largest statistically significant inverse association, cg07850633, was within the body of the MACRO domain-containing protein 2 (*MACROD2*) gene. A SNP within *MACROD2* has been associated with autism spectrum disorder (ASD) (66), though other studies have yielded potentially contradictory evidence (67). Even among persons without a diagnosis of intellectual disability or ASD, this same genetic variant is associated with autism-like traits (68). Additionally, *MACROD2* has been implicated in numerous other neurological disorders including attention-deficit/hyperactivity disorder (69), schizophrenia (70), and temporal lobe volume (71). While nonsense variants in *EBF3*, which encode for the early B Cell Factor 3, may contribute to developmental delay and intellectual impairment (72). Genetic variation in the Ral Guanine Nucleotide Dissociation Stimulator Like 1 (*RGL1*) gene, has been associated with attention (73) and conduct problems among children with attention deficit hyperactivity disorder (74). Additionally, genetic mutations within *CACNA1H*, which encodes for a subunit of a voltage gated calcium channel, lead to decreased calcium channel activity in neuronal cells, and have been linked to ASD (75), to epilepsy (76), and to amyotrophic lateral sclerosis (77). Overall, these findings suggest that our NNNS-associated epigenetic variations occurred at numerous genomic regions with recognized roles in neurodevelopmental or neurodegenerative disorders, which may provide a link between preterm birth and poorer neurodevelopmental outcomes.

There were some limitations to this study. We used a false discovery rate of 10% to identify significantly differentially methylated CpG sites, and thus it is probable that some of identified epigenetic loci are false-positives. We encourage additional investigation of infant DNAm, epigenetic age, and neurobehavior to determine whether similar relationships can be observed in independent populations. We utilized buccal cells as a surrogate tissue to examine the relationships between neurobehavioral profiles and DNAm, as it is not possible to perform such examinations in the neuronal tissues. However, for studies of children, buccal cell collection leads to greater compliance (78). Recent evidence also suggests that buccal samples may be very appropriate for epigenetic analyses of neurodevelopmental outcomes, as they arise from the same germ cell layer as the brain and thus may share similar early epigenetic patterning and susceptibility (79-81), and have demonstrated (DNAm) variability associated with later neurobehavioral outcomes. Yet, we remain cautious in the interpretation of these observations in terms of mechanism. These data were collected and analyzed cross-sectionally, so we cannot infer directionality of the observed relationships between NNNS profiles with DNAm or epigenetic age. It is notable, however, that the atypical profile observed by us and others in different populations, is related to epigenetic markers including with preterm infants and predicts long term developmental outcome. Thus, it is possible that the combination of epigenetic changes and NNNS profiles may lead to the early identification of which individual children are most at risk for adverse developmental outcome. Longitudinal studies of epigenomics and neurobehavioral outcomes are needed to establish whether epigenetic variations are detectable prior to the presentation of neurobehavioral impairments, and to examine whether and how these potential predictors vary throughout early life development.

## Conclusions

We found that among very preterm infants (< 30 weeks PMA), those with an optimal neurobehavioral profile had slightly older epigenetic age, while infants with an atypical neurobehavioral profile had differentially methylated CpGs at multiple genes linked to neural structure, function, or different neurobehavioral or neurodegenerative conditions. These relationships were detected using buccal cell DNAm, building upon the existing evidence that buccal cells may be a suitable surrogate tissue for studying neurobehavioral conditions in human observational studies. Three of the NNNS-associated CpGs were within genes yielding multiple levels of evidence for plausible roles in neurobehavioral health, annotated to *PLA2G4E*, *TRIM9*, and *GRIK3*, all of which were among the significantly enriched GO-terms or neuronal pathways, and linked to neurobehavioral disorders. The combination of epigenomics and neurobehavior holds promise for a personalized medicine approach to the early detection of children most at risk for poor developmental outcome.

---

**Supplemental Figure 1:**
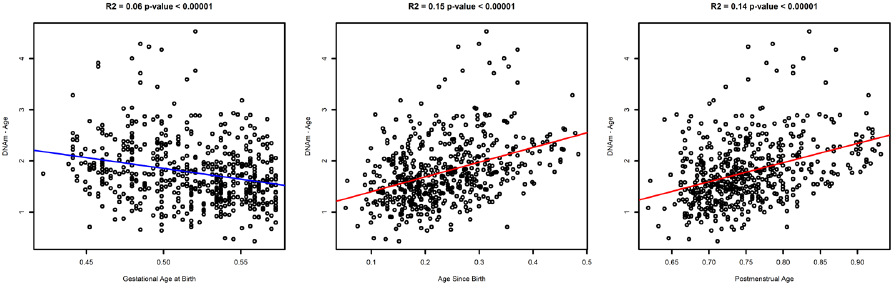
Relationships between epigenetic age with gestational age, chronological age since birth, and postmenstrual age (ages are represented as days divided by 365).

**Supplemental Figure 2:**
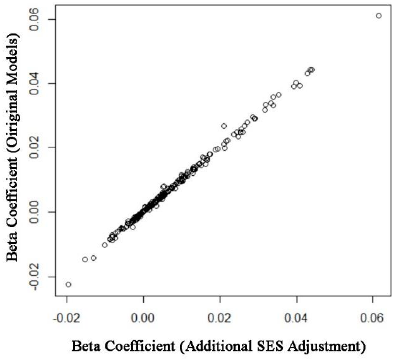
Comparison of beta-coefficients before and after adjustment for low SES, among the 250 CpGs that yielded the most statistically significant associations with the atypical NNNS profile.

